# Sex, head size, and male mate location behavior affect allometric scaling of the eyes in *Centris pallida* (Hymenoptera: Apidae) bees

**DOI:** 10.64898/2026.01.09.698656

**Authors:** M Barrett, S O’Donnell

## Abstract

Variation in eye structure and function can allow organisms to exploit novel environments or perform unique behaviors. However smaller body sizes may impose morphological constraints on eye investment, limiting adaptive behavioral performance. Males of the solitary desert bee *Centris pallida* (Hymenoptera: Apidae) are dimorphic in both body size and sensory mate location behaviors: large-morph males are fixed on a scent-based strategy while small-morph males generally use a sight-based strategy. Females are not known to vary categorically in mating behavior but span nearly the full range of male body sizes, allowing for sex comparisons in allometric scaling. We hypothesized that sex, body size, and male mate location behavior would impact eye structure and estimated global and localized visual acuity in *C. pallida* bees. We predicted that male and female body size would correlate with ommatidia number and size, and visual acuity, due to allometric constraints. We further predicted that small-morph males would have increased relative investment in the eyes and improved visual acuity compared to large-morph males. We found that males had increased ommatidia numbers and larger eyes than similarly-sized females, as well as greater average and maximum ommatidia diameters. Females had improved localized visual acuity in the dorsofrontal hotspot than small-morph males, but similar acuity to large-morph males. Body size thus impacted localized acuity in the dorsofrontal region in males but not in females. Small-morph males had increased relative eye size and ommatidia density but reduced total ommatidia numbers, eye surface areas, and estimated visual acuities compared to large-morph males.

## Introduction

Bees use visual cues to find mates, navigate a complex 3D environment, locate resources or nests, learn multimodal associations, and defend themselves or their nests from predation or parasitism (Streinzer & Spaethe 2014, Reinhard et al. 2004, Srinivasan 2010, Guédot et al. 2005, Inouye 2000, Collet 1992, Breed et al. 2004, Wittman 1985, Wittman et al. 1990). Size and functional capacity of bees’ eyes often evolve to match ecological and behavioral challenges. For example, crepuscular or nocturnal carpenter bee species often have unique anatomical features that enhance light sensitivity compared to similarly-sized, closely-related but diurnally foraging species (Somanathan et al. 2008). Male bumble bees that rely on visual perching strategies to find mates have enlarged eyes with higher spatial resolution and a frontal region with greater light sensitivity relative to males of species that patrol scent routes (Streinzer & Spaethe 2014). Specialization in external sensory structures can thus support the exploitation of novel environments or performance of unique behaviors.

Here we explore the possibility that eye structure and function exhibit behavior-related adaptive variation within a bee species. Males of the solitary desert bee *Centris pallida* (Hymenoptera: Apidae) are dimorphic in both body size and sensory mate location behaviors: large-morph males are fixed on a scent-based mate location strategy, while small-morph males generally use a sight-based mate location strategy (though they are behaviorally flexible; Snelling 1984, Alcock 1976, Alcock et al. 1976, Alcock et al. 1977, Alcock 1984). There is neuroanatomical specialization associated with these strategies: large-morph males have increased relative investment in brain tissue devoted to chemosensory information (antennal lobes) while small-morph males have increased relative investment in tissue devoted to visual information (optic lobes; Barrett et al. 2021). Further, small-morph males have reduced cell densities in their optic lobes, suggesting that specialization occurs at multiple levels of sensory organization in male *C. pallida* (Barrett and Godfrey, *in review*).

Unlike brain tissue investment, which is plastic and may shift in response to *or* expectation of behavior (Baudier et al. 2021, O’Donnell et al. 2021), external visual anatomy is always fixed in bees following metamorphosis and thus represents investment and constraints in sensory capabilities modalities prior to their use. The external visual system of bees mediates the input of visual cues and consists of paired apposition compound eyes, made up of repetitive units called ommatidia that receive light through small lens to form an image that is a composite of the spots generated by each ommatidium. Each ommatidium thus represents a point sampled in space, and therefore the number of ommatidia, as well as inter-ommatidial angle, generally correlate with spatial resolution and visual acuity (Land 1997, Cronin et al. 2014). Ommatidium size is also important, with larger facet diameters bringing in more photons of light and thus increasing light sensitivity. Comparative studies suggest body size can constrain bee visual systems, with bees experiencing tradeoffs between ommatidia number and size; body size is a major predictor of the number and diameter of ommatidia across sbee species (Jander & Jander 2002). To maintain critical adaptive functions, some bee eyes show differential scaling across the eye surface that maximizes ommatidia diameters and localized acuity in critical visual regions while still preserving overall ommatidia number ().

Smaller bees (across or within species) generally have reduced eye surface areas that necessarily limit the global resolution and/or sensitivity of their eyes (Spaethe & Chittka 2003; Jander & Jander 2002). For instance, smaller-bodied stingless bee species have reduced ommatidia diameters, decreasing light sensitivity and limiting foraging in dim-light conditions (Streinzer et al. 2016). Within species, larger-bodied bumblebee workers have increased light sensitivity, image resolution, and target detection abilities compared to smaller-bodied workers, as both ommatidia diameter and number correlate positively with body size (Maebe et al. 2015, Spaethe & Chittka 2003, Kapustjankskij et al. 2007; though see: Kapustjanskij et al. 2007, Streinzer & Spaethe 2013). However, for females of solitary species like *C. pallida*, all individuals must engage in every task in order to be reproductively successful; therefore, body size may place different constraints on the evolution of the visual system in size-variable solitary female bees, compared to social bees.

We hypothesized that allometric constraints associated with sex and body size, as well as male mate location behavior, would correlate with eye structure and expected function in *C. pallida* male and female bees. We predicted that, within-morph (for males) and in females, smaller body sizes would correlate with reduced numbers of ommatidia, decreased average ommatidia size, and reduced visual acuity, due to the allometric constraints of reduced body size. We further predicted that small-morph males would have increased relative investment in the eyes and improved visual acuity, due to their use of a visual mate location strategy.

## Materials and Methods

### Specimen collection

Adult *C. pallida* males and females were collected in late April and early May of 2018 in Arizona (N33.552°, W-111.566°) where dense emergence and nesting aggregations have persisted for several decades (e.g. Alcock et al. 1976). Heads were cut from the thorax and placed immediately into Prefer fixative (Anatech, Ltd.) following weighing to the nearest 0.1 mg on an analytical balance. Males were classified as large or small-morph males using a combination of behavior and morphology, as in a previous study (Barrett et al. 2021). Frontal photographs of the head capsules, and dorsal photographs of the thorax, under a dissecting scope were used to obtain head height, eye height, head width (Alcock et al. 1977), and intertegular span (Cane 1987).

### Eye casts and ommatidia counts

Eye casts and ommatidia counts were generally conducted as in Streinzer et al. (2013). Briefly, bee heads (n = 18 males, 8 females) were dried overnight before being mounted on a pyramid made of clay. A thin layer of Sally Hansen 101 top coat nail polish was applied to the left eye and allowed to partially dry, then carefully removed from the eye using #5 forceps (Fine Science Tools); if the left eye was damaged, the right eye was used instead. The interior surface of the cast was placed facing upward on a small drop of distilled water, on gelled (gelatin from bovine skin, chromium potassium sulfate) slides. Cuts were made in the cast using microscissors to allow the cast to flatten on the slide, and the water was allowed to evaporate from underneath the cast at room temperature (inside a box to minimize particulate contamination).

Casts were photographed at 10X magnification using a compound light microscope-mounted digital camera with a 1 X camera mount. Digital photographs were taken using LAS V4.9 software, with sharpness set to robust, and LAS was used to automatically stitch several photographs together into a complete image of each cast. A photograph of a micrometer at 10X was used to convert pixels to mm. The FIJI (Schindelin et al. 2012) freehand selection tool was used to trace the outer surface of the cast, in order to obtain the surface area of the eye.

The FIJI multipoint tool was used to manually select the center of each ommatidium (n = 8 casts for females and large males, n = 10 casts for small males), also providing the number of ommatidia. The XY coordinates of these points were uploaded to Meshlab (Visual Computing Lab – ISRI – CNR, http://meshlab.sourceforge.net/) in order to obtain ommatidia diameters (estimated using the distance between neighboring ommatidia diameters). Inkscape (https://inkscape.org/) was used to portray the changing pattern of ommatidia diameters across the eye surface, using a color gradient.

### Inter-ommatidial angle, eye curvature, and visual acuity estimates

Global inter-ommatidial angle (ΔΦ_G_) was estimated using Land (1997)’s formula, which assumes a hemispheric visual field (where n is the total number of ommatidia):

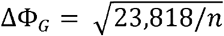

Regional patterns of ommatidia diameter and curvature may allow for variable ΔΦ across the eye surface (Scales & Butler 2015); therefore, the radius of curvature method (Bergman & Rutkowski 2016) was used to obtain localized inter-ommatidial angle along the x-axis (ΔΦ_x_; parallel to horizon; measures were taken 20° from inner edge of eye, viewed dorsally) and y-axis (ΔΦ_y_; perpendicular to horizon; measures were taken 20° dorsally from perpendicular bisection of the eye, viewed laterally) in the dorsofrontal ‘hot spot’, using photos of the head capsule taken on a dissecting scope in the dorsal and lateral orientation, respectively. FIJI was used as in Duncan et al. (2021) to calculate localized eye curvature along each axis (Figure S1; distance, *b*, of eye surface covered in a given angle, *a*). The average diameter of five facets in each direction (*D*) was calculated using facets from the flattest region of the eye near where the curvature was measured, using photographs taken in the frontal orientation. Inter-ommatidial angle was calculated as:

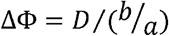

Visual acuity was estimated as twice the ΔΦ, as in Land (1997).

### Statistical analyses

GraphPad Prism v. 9.1.2 (GraphPad Prism for Windows 2021) was used for all statistical analyses and data were confirmed to meet the assumptions of parametric tests before those analyses were performed using Anderson-Darling and Shapiro-Wilk normality tests and an F-test was used to test for equal variance. Alpha was set to 0.05 for all analyses. Unpaired t-tests (between male morphs) or ANOVAS (between male morphs and females) were used to analyze differences in head width, IT span, body mass, ommatidia number and density, largest and smallest ommatidia diameter, localized eye curvature in the x and y axis, and ΔΦ_G_, ΔΦ_x_ and ΔΦ_y_ between large and small males. As head size seemed most relevant to eye allometry, and is the measure commonly used when documenting male morph body size in this species (Alcock et al. 1989, 1995, 2013, Barrett and Johnson 2022, Barrett 2025), we used head width as the measure of body size in all analyses where head width was not the dependent variable.

Linear regressions were used to analyze relationships between log(head width) and log(intertegular span), log(eye height) and log(sqrt(eye surface area)), log(head width) and log(eye height), head width and relative eye height, ommatidia number, ΔΦ_x,_ and ΔΦ_y_, and male relative optic lobe investment with global and averaged localized visual acuity.

## Results

### Body Size

Small- and large-morph *C. pallida* males differed in mean head width, IT span, and wet body mass (Figure 1: unpaired t-tests; head width: t = 8.84, df = 28, p < 0.0001; IT span: t = 5.36, df = 21, p < 0.0001; body mass: t = 11.44, df = 27, p <0.0001). There was no overlap in head width or body mass: large-morph males had a minimum head with of 5.34 mm and body mass of 0.25 g (ranges, head width: 5.34 – 6.02 mm; body mass: 0.25 – 0.35 g) while small-morph males had a maximum head width of 5.27 mm and body mass of 0.21 g (ranges, head width: 4.53 – 5.27 mm; body mass: 0.11 – 0.21 g). Large and small-morph males overlapped slightly in IT span, with a minimum IT span of 5.00 mm for large-morph males (range: 5.00 – 5.77 mm) and a maximum IT span of 5.11 mm for small-morph males (range: 4.11 – 5.12 mm).

**Figure 1.**
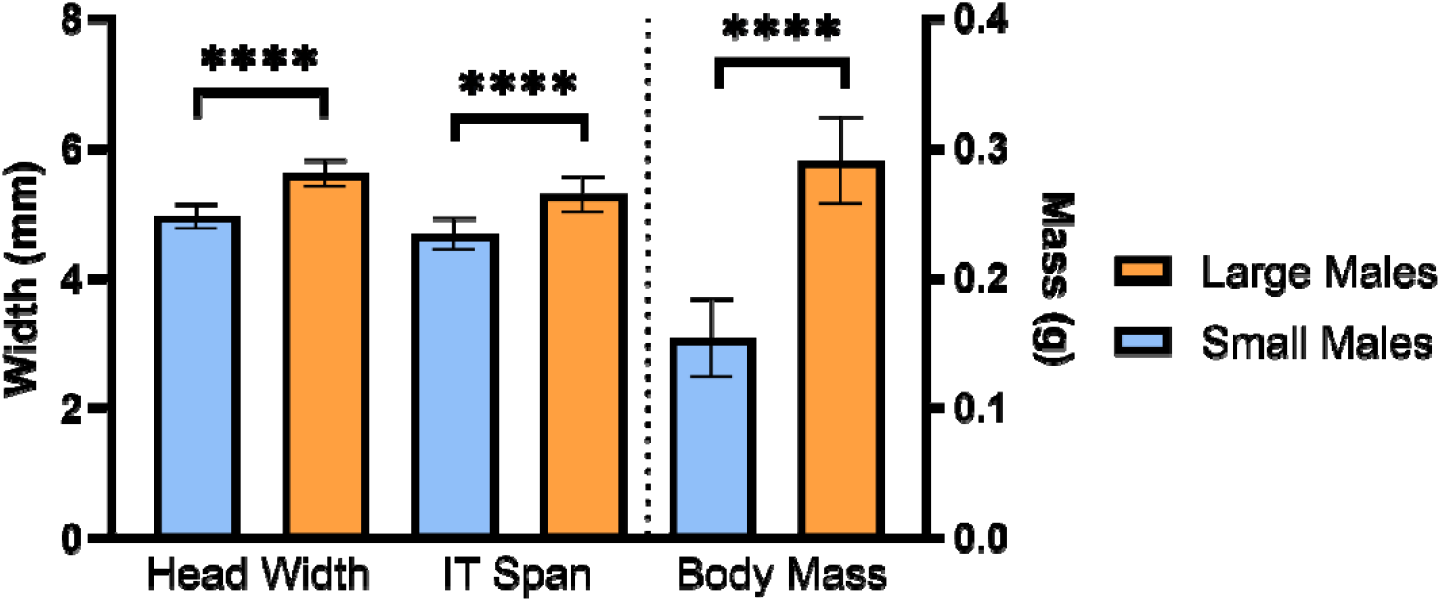
Differences in body size between large and small-morph males. Small- and large-morph *C. pallida* males differed in mean head width, IT span, and wet body mass. **** = p<0.0001

Male morphs did not differ in the scaling of head size to intertegular span (F = 0.005, df = 19, p = 0.94), so all males were plotted together. Head width correlated hypoallometrically with intertegular span in both males and females, suggesting smaller individuals have relatively increased head capsule sizes (Figure S2: linear regression, males: log [head width] = 0.75 log [Intertegular span] + 0.20, R^2^ = 0.82, F = 96.73, n = 23, p < 0.0001; females: [head width] = 0.81 [Intertegular span] + 0.19, R^2^ = 0.71, F = 26.82, n = 13, p = 0.0003).

Females overlapped extensively with males, ranging from 0.16 to 0.35 g in mass, with IT spans of 4.17 to 5.25 mm, and head widths of 4.85 – 5.97 mm.

### Eye Size

Eye surface area differed between large and small-morph males (Unpaired t-test; t = 4.79, df = 16, p = 0.0002), with a mean of 9.49 ± 0.51 mm^2^ in large males and 8.21 ± 0.60 mm^2^ in small males. Females had smaller mean eye surface areas than both large and small morph males (ANOVA, F = 64.19, df = 23, p < 0.0001; Bonferroni-correct MCT, SM-F: t = 7.03, df = 23, p < 0.0001; LM-F: t = 11.26, df = 23, p < 0.0001), with a mean of 6.35 mm^2^ (range: 5.48 – 7.19 mm^2^).

Large and small-morph males differed significantly in the allometric scaling of their eye height (Figure 2A; test for difference in the regression slopes: F = 9.55, df = 26, p = 0.0047). Small-male eye height scaled hyperallometrically (*b* = 1.10; linear regression with head width, R^2^ = 0.90, F = 148.9, df = 16, p < 0.0001) while large-male and female eye heights scaled hypoallometrically (large males: *b* = 0.61; R^2^ = 0.67, F = 20.18, df = 10, p = 0.0012; females: *b* = 0.83; linear regression, R^2^ = 0.84, F = 55.94, df = 11, p < 0.0001).

**Figure 2.**
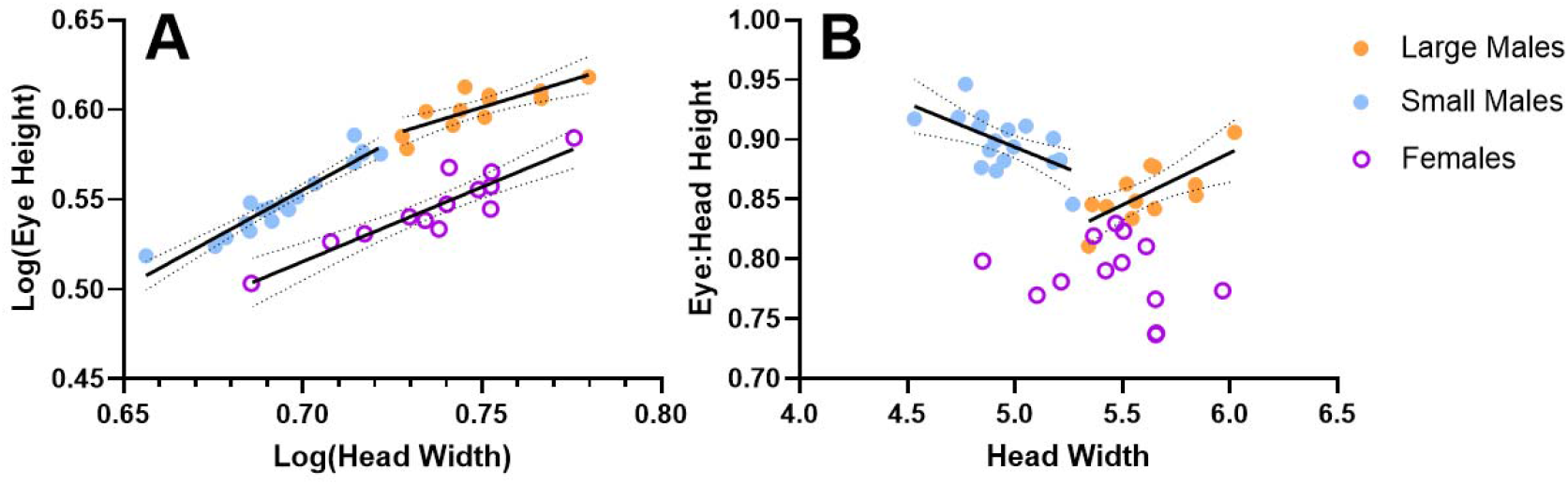
Small morph male eye heights scale hyperallometrically (*b* > 1.0), while large morph male and female eye heights scale hypoallometrically (*b* < 1.0). A) Scatterplot showing that large and small-morph males differed significantly in the allometric scaling of their eye heights. Small-morph male eye heights scaled hyperallometrically (b = 1.10) while large-morph male eye heights scaled hypoallometrically (b = 0.61), as did female eye heights (b = 0.83). B) Scatterplot showing that small-morph males had increased relative eye heights at smaller body sizes, while large-morph males had decreased relative eye heights at smaller body sizes; there was no relationship in females. Solid lines = significant linear regression; dotted lines = 95% CI

Relative eye:head height also differed as a function of body size in male morphs (Figure 2B; test for differences in regression slopes: F = 21.53, df = 26, p < 0.0001). Small males had increased relative eye heights at smaller body sizes (linear regression, [Rel eye:head height] = −0.07 [head width] + 1.26, R^2^ = 0.39, F = 10.37, df = 16, p = 0.0054), while large males had decreased relative eye height at smaller body sizes ([Rel eye:head height] = −0.09 [head width] + 0.37, R^2^ = 0.54, F = 11.61, df = 10, p = 0.0067). Relative eye height did not change as a function of body size in females (linear regression, R^2^ = 0.07, F = 0.81, df = 11, p = 0.39); females had reduced relative eye heights compared to both male morphs (ANOVA, F = 69.08, df = 40, p < 0.0001; Bonferroni MCT, F-LM: t = 6.65, p <0.0001; F-SM: t = 11.74, p < 0.0001).

### Ommatidia Number, Diameter, and Distribution

Large males had more ommatidia than small males (Unpaired t-test: t = 2.67, df = 16, p = 0.0166). Ommatidia number increased with increasing body size at a similar rate in both males and females (Figure 3A; males, linear regression: [Ommatidia number] = 1095 [Head width] + 5198, R^2^ = 0.56, F = 19.96, df = 16, p = 0.0004; females, linear regression: [Ommatidia number] = 1247 [Head width] + 3651, R^2^ = 0.75, F = 18.36, df = 6, p = 0.0052; test for difference between slopes, F = 0.10, df = 22, p = 0.75); however males had more ommatidia than females of the same body size (test for differences between intercepts, F = 18.85, df = 23, p = 0.0003).

**Figure 3.**
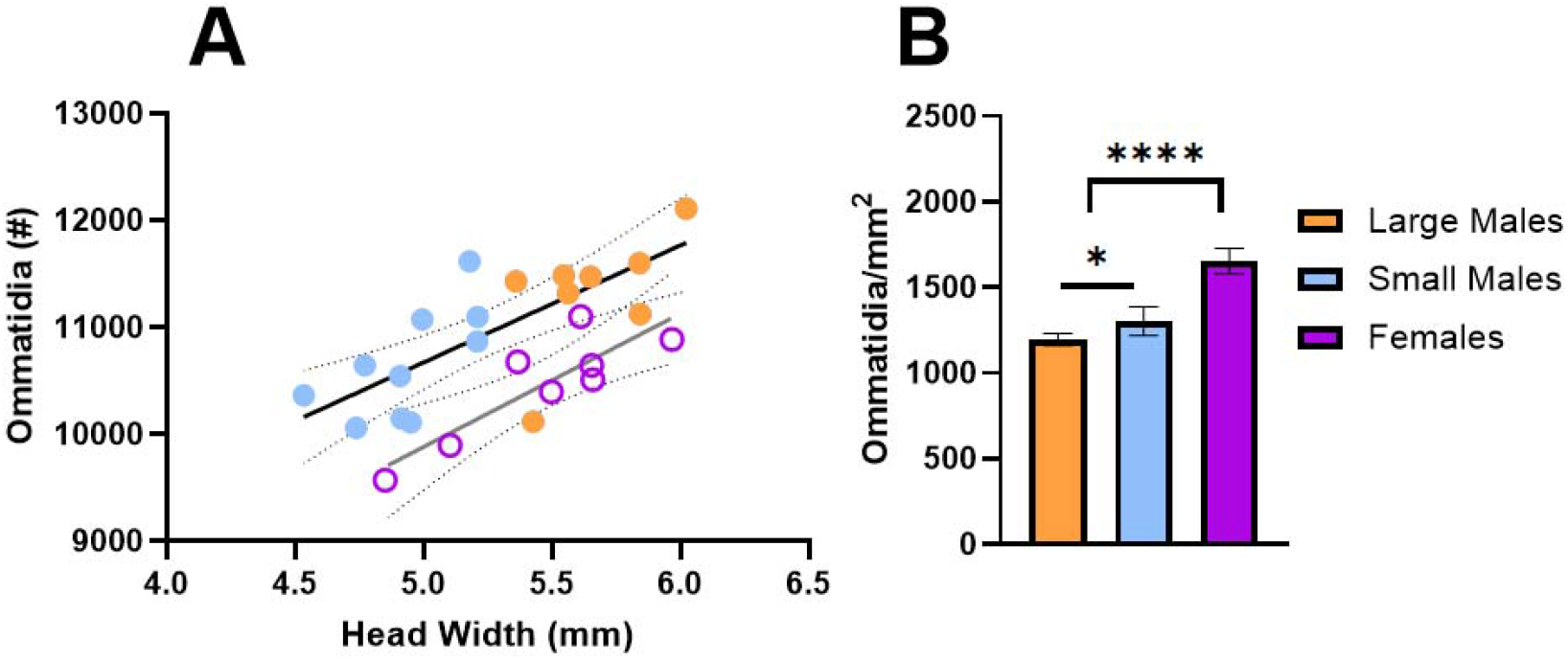
Ommatidia number and density in males and females. A) Scatterplot showing ommatidia number increased with increasing body size at a similar rate in both males and females; however, males had more ommatidia than females of the same body size. Solid lines = significant linear regressions; dotted lines = 95% CI. B) Bar chart showing the density of ommatidia varied in male morphs and females. * = p < 0.05; **** = p < 0.0001

The number of ommatidia per mm^2^ of eye surface area was lower in large compared to small-morph males, and in both male morphs compared to females (Figure 3B: One-way ANOVA, F = 95.61, df = 23, p < 0.0001). Small males had a lower proportion of ommatidia above 28 μm diameter, and higher densities of ommatidia below 26 μm diameter (particularly in the 18 − 26 μm diameter range; Figure 4). Large and small-morph males did not differ in the size of their largest ommatidia (Table 3-1; ANOVA, Bonferroni MCT: t = 1.16, df = 23, p = 0.78), but small-morph males had smaller average ommatidia diameters and also had the smallest ommatidia (average: t = 3.31, p = 0.0095; smallest diameter: t = 3.62, p = 0.0043). Females had a narrower range of ommatidia diameters, with a greater proportion of ommatidia between 18 and 26 μm compared to both large and small males (Figure 4) and reduced averaged and largest ommatidia diameters (Table 1).

**Table 1.** Mean ± SD (range) largest, smallest, and average ommatidia diameters of small and large males, and females.

|  | Small-morph males | Large-morph males | Females |
| --- | --- | --- | --- |
| Smallest diameter | 15.30 ± 0.86<sup>a</sup><br>(13.82 – 16.34) | 17.08 ± 0.85<sup>b</sup><br>(16.08 – 18.53) | 15.91 ± 1.38<sup>a,b</sup><br>(14.09 – 18.51) |
| Average diameter | 29.25 ± 0.95<sup>b</sup><br>(27.96 – 30.92) | 30.37 ± 0.42<sup>c</sup><br>(29.80 – 31.19) | 26.05 ± 0.55<sup>a</sup><br>(25.39 – 27.18) |
| Largest diameter | 46.19 ± 2.45<sup>b</sup><br>(41.52 – 49.38) | 47.27 ± 1.36<sup>b</sup><br>(45.49 – 49.91) | 34.13 ± 1.78<sup>a</sup><br>(31.98 – 36.84) |

<sup>a,b,c</sup> letters indicate significant differences between morphs/females in that ommatidia size category (e.g., row; One-way ANOVA, Bonferroni MCT; all $p < 0.01$ ).

**Figure 4.**
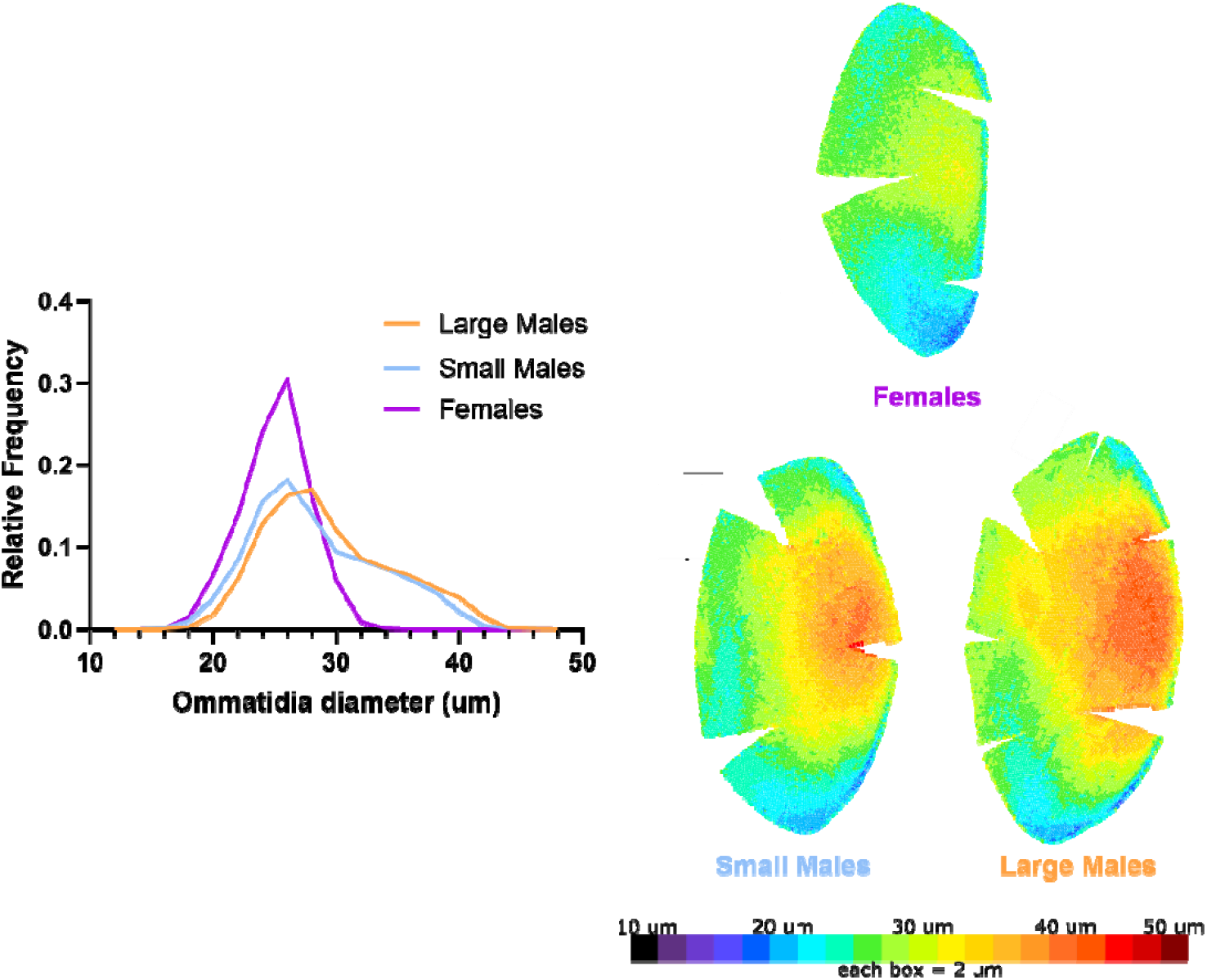
Relative frequency of ommatidia diameters varies in small and large-morph males and females. A representative, mirrored cast of the left eyes of a small and large-morph male, and a female, shows the hot-spot of largest-diameter cells, dorsofrontally (scale bar, upper left of small male cast = 0.5 mm; to see all casts, Figures S3 and S4).

Large and small males had a dorsofrontal “hot spot” with their largest diameter ommatidia (Figure S3). In females, there was still a bias towards larger ommatidia diameters in the dorsofrontal region, but the largest ommatidia in that region were still smaller than in males (p < 0.01; Table 1; Figure S4).

### Inter-ommatidial angle, eye curvature, and visual acuity estimates

Global inter-ommatidial angles (ΔΦ_G_) based only on ommatidia number (not accounting for possible differences in curvature) were slightly, but significantly, greater in small-morph males compared to large-morph males (Table 2, Figure 5; ANOVA, Bonferroni: t = 2.61, df = 23, p = 0.047). Females had larger ΔΦ_G_ than large-morph but not small-morph males (LM-Fem: t = 3.22, p = 0.0144; SM-Fem: t = 0.79, p > 0.99).

**Table 2.** Mean ± SD (range) of local ΔΦ_x_, ΔΦ_y_, and global ΔΦ_G_ inter-ommatidial angles in small and large males and females.

|  | Small-morph males | Large-morph males | Females |
| --- | --- | --- | --- |
| $\Delta\Phi_x$ | $2.28^\circ \pm 0.38$ ($1.47^\circ - 2.94^\circ$)<sup>a</sup> | $1.56^\circ \pm 0.24^\circ$ ($1.18^\circ - 1.84^\circ$)<sup>b</sup> | $1.72^\circ \pm 0.26^\circ$ ($1.39^\circ - 2.27^\circ$)<sup>b</sup> |
| $\Delta\Phi_y$ | $1.30^\circ \pm 0.28$ ($0.71^\circ - 1.80^\circ$)<sup>a</sup> | $0.94^\circ \pm 0.20^\circ$ ($0.63^\circ - 1.30^\circ$)<sup>b</sup> | $0.80^\circ \pm 0.16^\circ$ ($0.56^\circ - 1.01^\circ$)<sup>b</sup> |
| $\Delta\Phi_G$ | $1.50^\circ \pm 0.04$ ($1.43^\circ - 1.54^\circ$)<sup>a</sup> | $1.45^\circ \pm 0.04$ ($1.40^\circ - 1.53^\circ$)<sup>b</sup> | $31.51^\circ \pm 0.04$ ($1.47^\circ - 1.58^\circ$)<sup>a</sup> |

<sup>a,b,c</sup> letters indicate significant differences between morphs/females in that ommatidia size category (e.g., row; One-way ANOVA, Bonferroni MCT; all $p < 0.01$ ).

**Figure 5.**
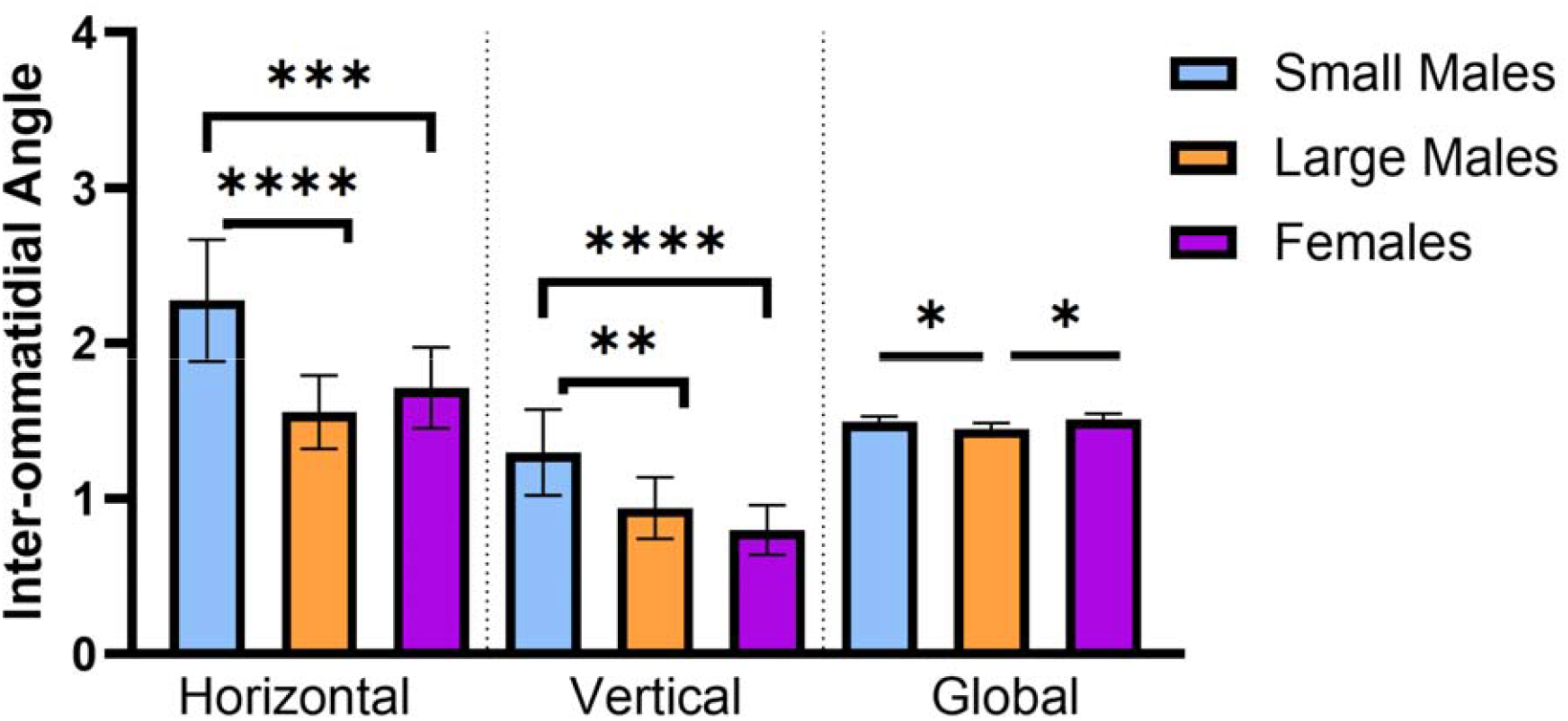
Small males have increased ΔΦ_x_ (horizonal), ΔΦ_y_ (vertical), and ΔΦ_G_ (global) compared to large males, and increased ΔΦ_x_ and ΔΦ_y_ compared to females. * = p < 0.05; ** = p < 0.01; *** = p < 0.001; **** = p < 0.0001

Using the RCE method to analyze localized inter-ommatidial angles in the dorsofrontal ‘hot spot’, small-morph males had increased ΔΦ_x_ (horizontal) and ΔΦ_y_ (vertical), and thus worse visual acuity, compared to large-morph males and females (Table 2; Figure 5; ANOVA, ΔΦ_x_: F = 18.52, df _x_ = 34, p < 0.0001; ΔΦ_y_: F = 16.80, df = 34, p < 0.0001). Large-morph males and females did not differ significantly from one another in either axis (all p > 0.05). In males, increased body size correlated with smaller interommatidial angles and thereby with improved visual acuity (Figure 6; linear regression: [ΔΦ_x_] = −0.93 [head width] + 6.84, R^2^ = 0.52, F = 27.2, df = 25, p < 0.0001; ANOVA of ΔΦ_y_: t = 3.56, df = 25, p = 0.0015; linear regression: [ΔΦ_y_] = −0.58 [head width] + 4.18, R^2^ = 0.53, F =27.82, df = 25, p < 0.0001); however, this was not the case in females (linear regression, x-axis: F = 0.01, df = 8, p = 0.92; y-axis: F = 0.06, df = 8, p = 0.82).

**Figure 6.**
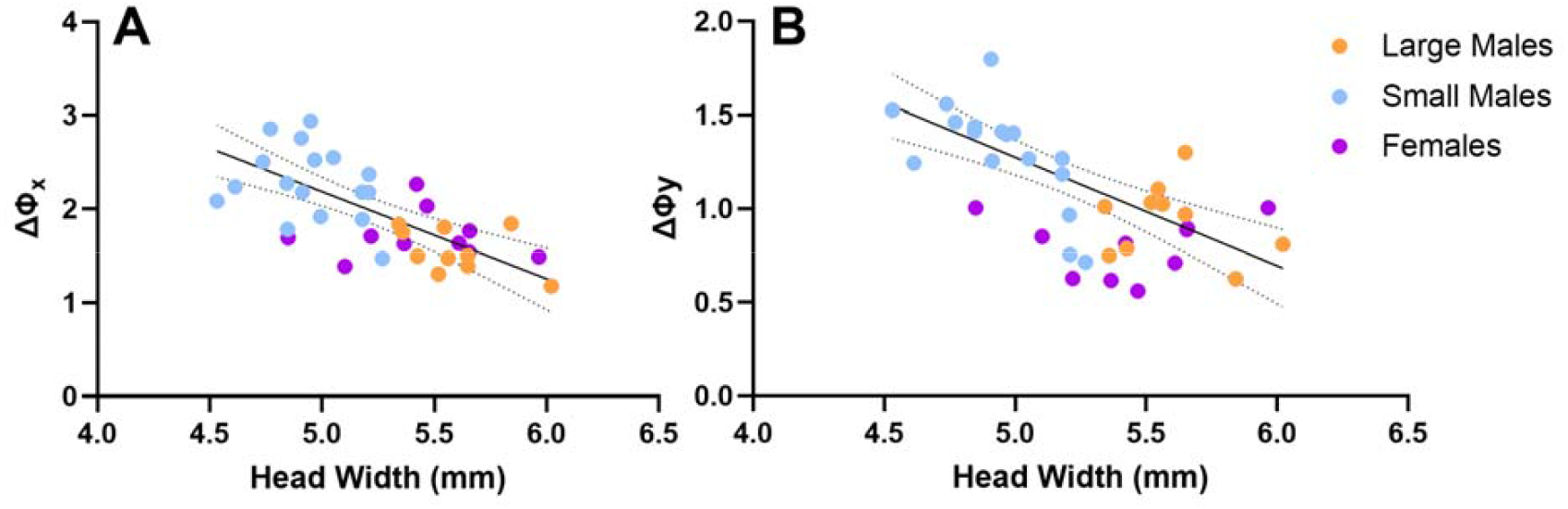
Head width negatively correlates with ΔΦ_x_ and ΔΦ_y_ in males but not females. Scatterplots showing increased body size correlates with decreased male A) ΔΦx and B) ΔΦy. Solid line shows significant linear regression for males; dotted lines = 95% CI.

Eye curvature and facet size combined to create these differences in visual acuity/inter-ommatidial angle across morphs and sexes (Kruskal Wallis or ANOVA, eye curvature: x-axis: K-W = 20.05, p < 0.0001, y-axis: F = 11.71, df = 34, p < 0.0001; facet diameter: F = 148.5, df = 34, p < 0.0001; y axis: F = 102.3, df = 34, p < 0.0001). Large males had reduced eye curvature (Figure S5A; Dunn’s or Tukey’s MCT; x-axis: z = 3.72, p = 0.0006; y axis: q = 6.23, df = 34, p = 0.0003) and increased facet diameters (Figure S5B; Tukey’s MCT, x axis: q = 4.78, df = 34, p = 0.0051; y axis: q = 4.71, df = 34, p = 0.0058) compared to small males. Large-morph males had reduced horizontal eye curvature compared to females (Figure S5A; Dunn’s MCT; x-axis, LM-Fem: z = 4.15, p < 0.0001) but similar vertical curvature (q = 1.14, p = 0.70). Small males had greater vertical eye curvature compared to females (q = 4.95, p = 0.0037) but similar horizontal curvature (z = 0.94, p > 0.99). Both large- and small-morph males had greater facet diameters compared to females (Figure S5B; Tukey’s MCT; x-axis, LM-Fem: q = 22.47, df = 34, p < 0.0001; SM-Fem: q = 20.43, df = 34, p < 0.0001; y-axis, LM-Fem: q = 18.93, df = 34, p < 0.0001; SM-Fem: q = 16.53, df = 34, p < 0.0001).

## Discussion

Sex, body size, and male mate location behaviors all correlate with differences in eye morphology and investment in *C. pallida* bees. The patterns of morph and sex variation suggest that some of these differences are adaptive, corresponding to behavioral challenges the bees face particularly in mating and courtship contexts.

Few studies have compared male and female bees of the same species, and the vast majority have compared social species where the impacts of external sensory systems on individual female fitness may be altered by colony-level selection and/or size-based division of labor. In *C. pallida*, we found eye size, average and largest ommatidia diameters, and ommatidia numbers were larger in males compared to females of a similar size. These data largely match categorical variation in means between the sexes in honey bees, orchid bees, and stingless bees (Streinzer et al. 2013, Ribi et al. 1989, Brand et al. 2018) though these species were not size-variable enough to provide allometric scaling data. In more size variable bumble bee species where allometric data could be obtained, only eye size and ommatidia diameters − but not ommatidia numbers − were larger in males compared to similarly-sized female workers (Kapustjanskij et al. 2007). While male *C. pallida* had increased ommatidia numbers compared to similarly-sized females, the effect of body size on ommatidia number did not differ between the sexes, similarly to bumble bees (Kapustjanskij et al. 2007).

Similar to other male bees that utilize a visual mate location strategy, male *C. pallida* irrespective of morph have an ‘acute zone’ of large-diameter ommatidia in the dorsofrontal region of the eye (Ribi et al. 1989, Streinzer et al. 2013). *C. pallida* males of both morphs had better visual acuity in the vertical axis over the horizontal axis in the acute zone, as did females. “Global” visual acuity was less variable between the morphs than localized visual acuity within the acute zone, suggesting that the changes in eye morphology that improve visual acuity at larger body sizes (in males) disproportionately benefit the dorsofrontal hotspot region. In size-variable female bumblebees, increased resources associated with their enlarged visual field can be invested unequally, with local increases in the resolution of the dorsofrontal region and global increases in light sensitivity (Taylor et al. 2019). Alongside disproportionately increased localized visual acuity in the acute zone, large-morph *C. pallida* males also have increased average ommatidia diameters and higher relative frequencies of larger-diameter ommatidia, suggesting global increases in light sensitivity. These intraspecific, allometric patterns in sensitivity and resolution may hold for bees irrespective of sex or sociality.

Small-morph *C. pallida* males, which primarily rely on a visual mate location strategy, had relatively increased eye:body size investment compared to large-morph males. There were also within-morph patterns in eye investment: small-morph male eyes scaled hyperallometrically, while large-morph male eyes scaled hypoallometrically. Thus, relative eye:head height investment differed between the morphs: smaller, small-morph males had relatively increased investment, while smaller, large-morph males had relatively decreased investment. This suggests that, at the smallest body sizes within a morph, eye size is differently prioritized. Interspecific studies of male bumblebees also show increased investment in the eyes when species primarily use visual, as compared to olfactory, mating strategies (Streinzer & Spaethe 2014), however our data show that intraspecific variation in mating behavior can also impact external sensory morphology.

Despite increased relative eye size, the reduced body sizes of small-morph *C. pallida* males still constrained their absolute eye size and its curvature and, thereby, their final visual acuity (as determined by localized interommatidial angle or total number of ommatidia) relative to large-morph males. Intraspecific studies in male bumblebees and size-variable honeybee drones have not quantified visual acuity, however there was no correlation between ommatidia number and body size in males of these species (Kapustjanskij et al. 2007, Streinzer & Spaethe 2013). In male *C. pallida*, body size still positively correlates with ommatidia number which, in combination with their worsened visual acuity, suggests the small males must ‘make the best of a bad situation’ in using the visual mate location strategy. As male *C. pallida* body sizes have declined over the last five decades (Barrett, 2025; Barrett and Johnson 2022), male bees may find their visual acuity and mating performance significantly disrupted by this shift towards smaller body sizes; tying eye allometry to functional changes in mating behavior could determine if species-level body size declines will impact fitness via changes in sensory morphology and ability.

Varying eye size, ommatidia numbers/diameters, and interommatidial angles allows for substantial variation in the visual-performance of individuals specializing on different behaviors within a species. Sex, body size, and male mating behaviors all correlated with variation in eye morphology, suggesting numerous variables can impact intraspecific external visual specialization. Despite increased relative investment in eye size for small-morph males, and morph-specific allometric scaling, small-morph males that rely primarily on a visual mate location strategy were unable to overcome the limitations of smaller body size in improving localized or global visual acuity and light sensitivity.

## Acknowledgments

We thank Stephen Buchmann for help collecting specimens in the field and Rachel Miller and Angelina Gomez for assistance with making casts and counting ommatidia. We thank Johannes Spaethe and, especially, Martin Streinzer for sharing detailed protocols from one of their papers on honey bee eye morphology which made this work possible.

## Supplementary Figures

**Figure S1.**
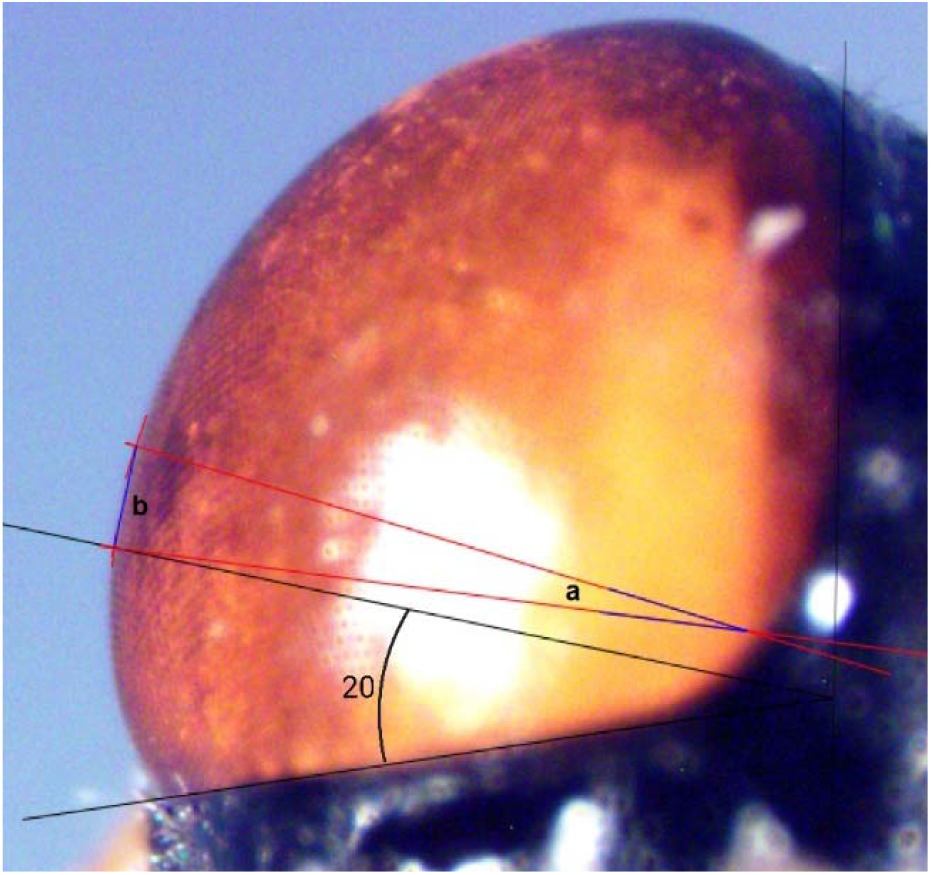
Radius of Curvature Estimation (RCE) of ΔΦx, 20° from the interior margin of the eye in the dorsal view. RCE method used to determine distance, *b*, of eye surface covered in a given angle, *a*, in both the dorsal (pictured) and lateral views.

**Figure S2.**
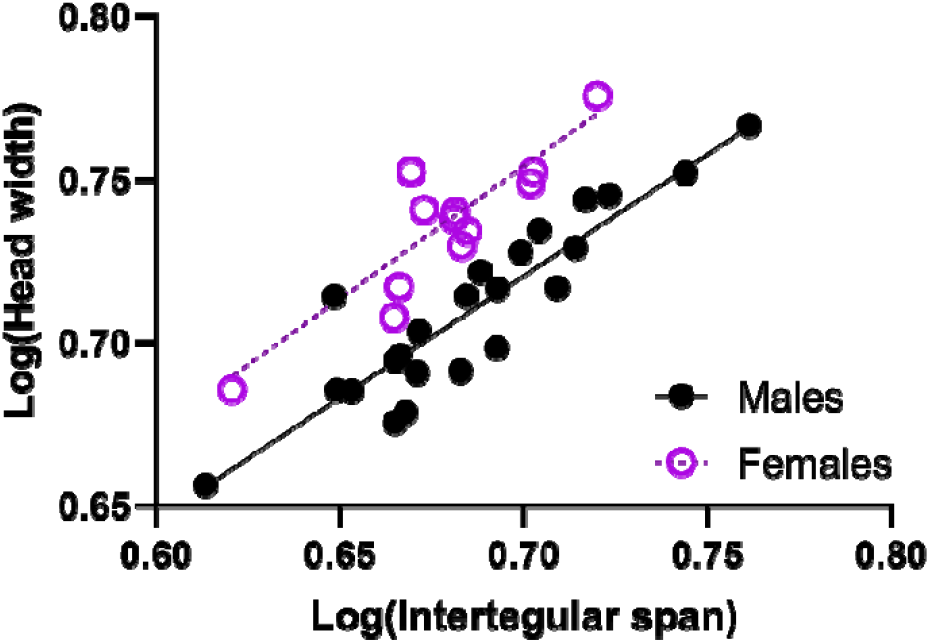
Head width positively correlates with intertegular span in both males and females. Head width correlates with intertegular span in males and females (linear regression, males: log [head width] = 0.75 log [Intertegular span] + 0.20, R^2^ = 0.82, F = 96.73, n = 23, p < 0.0001; females: [head width] = 0.81 [Intertegular span] + 0.19, R^2^ = 0.71, F = 26.89, n = 13, p = 0.0003).

**Figure S3.**
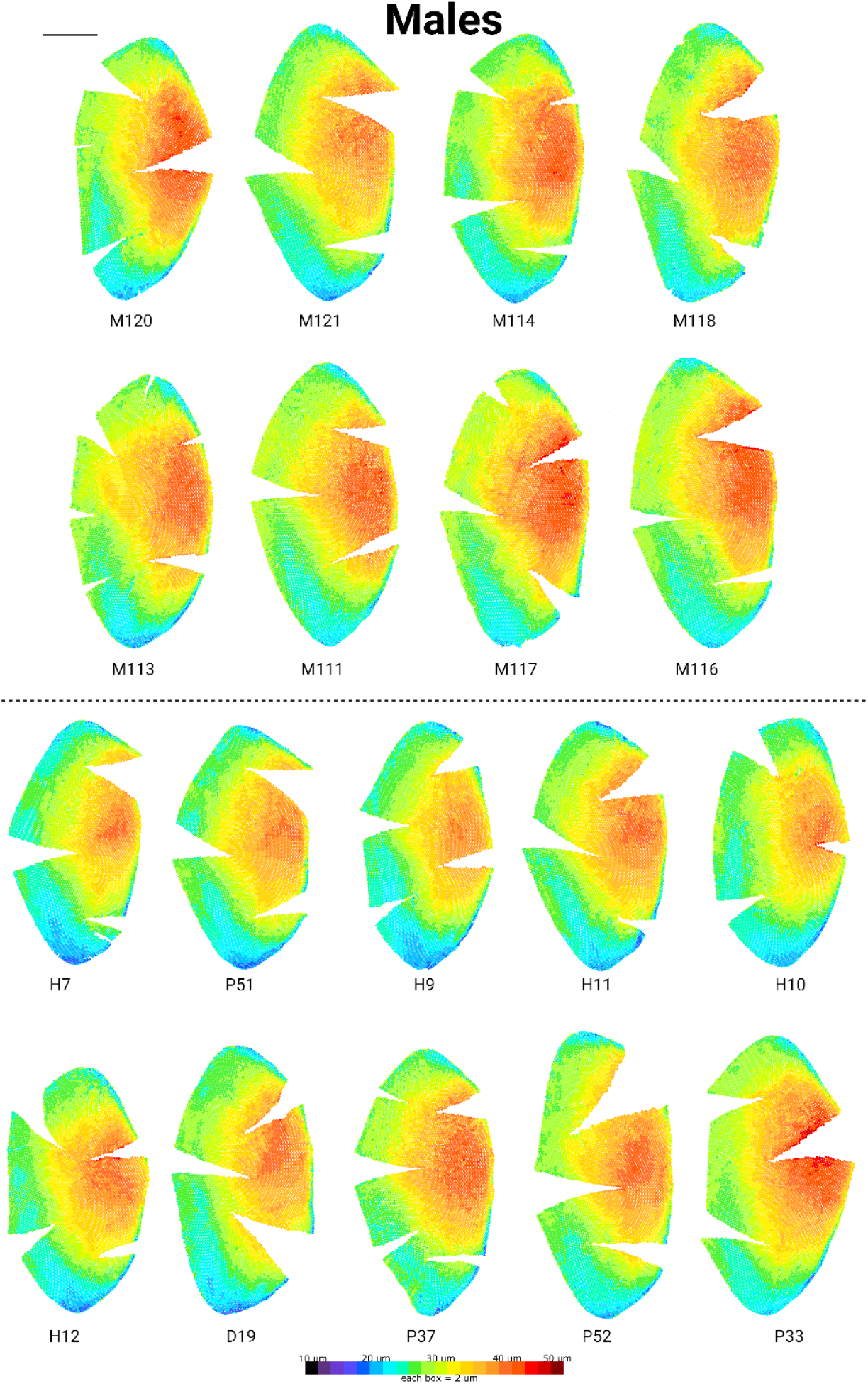
Casts of large (above dashed line) and small (below dashed line) *C. pallida* male morphs. Eye casts are arranged by increasing eye surface area within each morph (smallest, top left, to largest, bottom right). Black scale bar = 1 mm.

**Figure S4.**
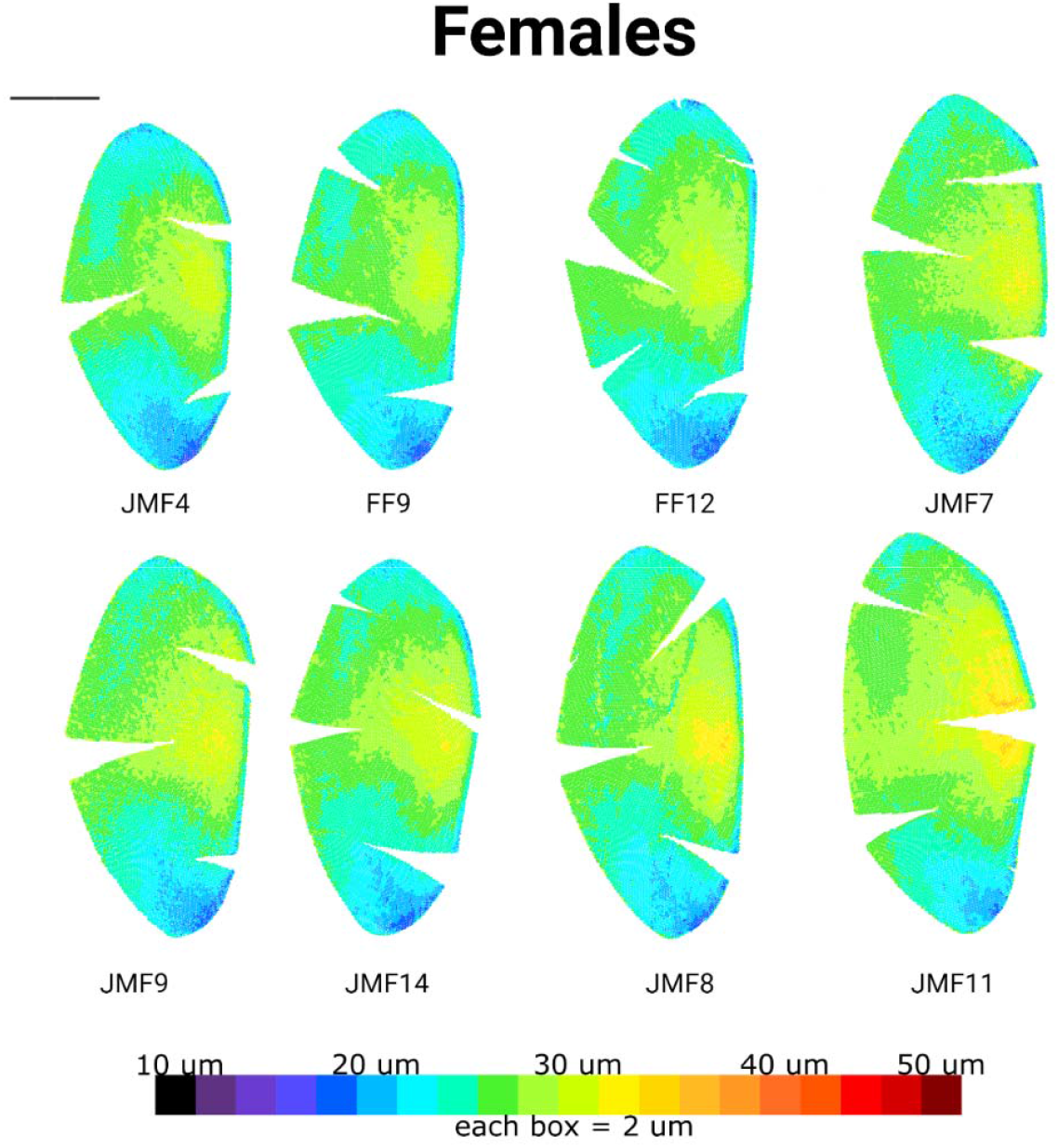
Casts of *C. pallida* females. Eye casts are arranged by increasing eye surface area (smallest, top left, to largest, bottom right). Black scale bar = 1 mm.

**Figure S5.**
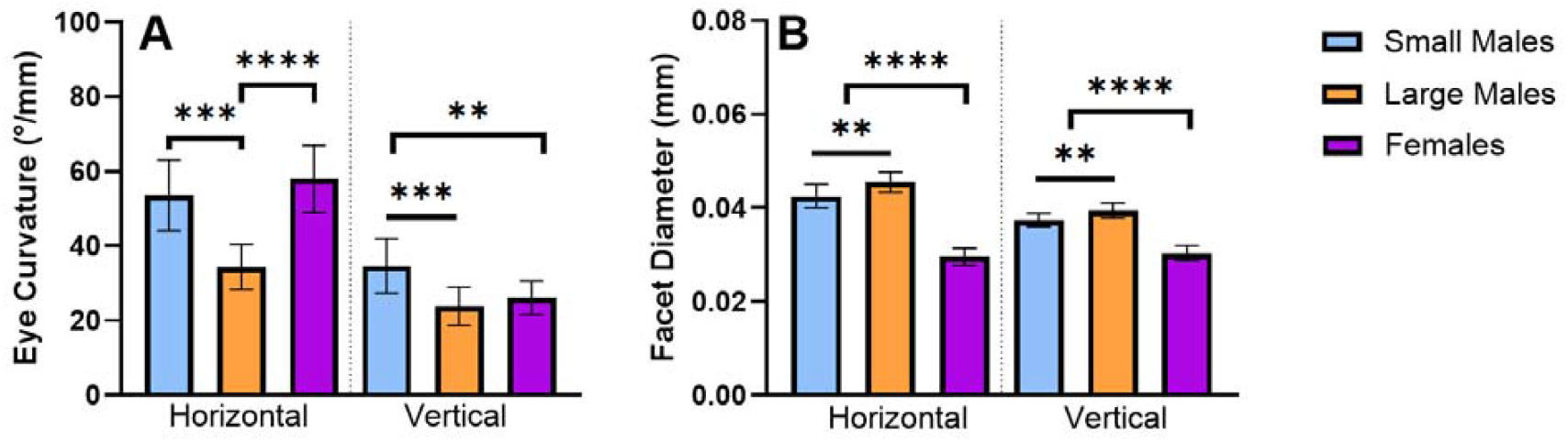
Changes in both eye curvature and facet diameter contribute to localized hotspot interommatidial variation in *C. pallida*. A) Large males have reduced eye curvature compared to small males; females have reduced vertical, but not horizontal eye curvature compared to small males. B) Females have smaller facet diameters than both large and small males; large males have increased facet diameters compared to small males. ** = p < 0.01; *** = p < 0.001; **** = p < 0.0001

